# Polylactic acid, a sustainable, biocompatible, transparent substrate material for Organ-On-Chip, and Microfluidic applications

**DOI:** 10.1101/647347

**Authors:** Alfredo E. Ongaro, Davide Di Giuseppe, Ali Kermanizadeh, Allende Miguelez Crespo, Arianna Mencatti, Lina Ghibelli, Vanessa Mancini, Krystian L. Wlodarczyk, Duncan P. Hand, Eugenio Martinelli, Vicki Stone, Nicola Howarth, Vincenzo La Carrubba, Virginia Pensabene, Maïwenn Kersaudy-Kerhoas

**Affiliations:** Institute of Biological Chemistry, Biophysics and Bioengineering, School of Engineering and Physical Science, Heriot-Watt University, Edinburgh EH14 4AS, United Kingdom; Division of Infection and Pathway Medicine, Edinburgh Medical School, College of Medicine and Veterinary Medicine, The University of Edinburgh, Edinburgh EH164SB, United Kingdom; Department of Engineering, Università degli Studi di Palermo, Viale delle Scienze building 5, 90128 Palermo, Italy; Department of Electronic Engineering, University of Rome Tor Vergata, Rome, Italy; School of Electronic and Electrical Engineering, Pollard Institute, University of Leeds, Woodhouse Lane, Leeds, LS2 9JT, United Kingdom; Institute of Photonics and Quantum Sciences, School of Engineering and Physical Science, Heriot-Watt University, Edinburgh EH14 4AS, United Kingdom; INSTM, Palermo Research Unit, Viale delle Scienze building 6, 90128 Palermo; ATeN Center, Università degli Studi di Palermo, Viale delle Scienze building 18, 90128 Palermo; School of Medicine, Leeds Institute of Medical Research, University of Leeds, Woodhouse Lane, Leeds, LS2 9JT, United Kingdom

**Keywords:** Organ-On-Chip, Poly-Lactic Acid (PLA), transparency, biocompatibility, rapid prototyping, cell tracking

## Abstract

Organ-on-chips are miniaturised devices aiming at replacing animal models for drug discovery, toxicology and studies of complex biological phenomena. The field of Organ-On-Chip has grown exponentially, and has led to the formation of companies providing commercial Organ-On-Chip devices. Yet, it may be surprising to learn that the majority of these commercial devices are made from Polydimethylsiloxane (PDMS), a silicone elastomer that is widely used in microfluidic prototyping, but which has been proven difficult to use in industrial settings and poses a number of challenges to experimentalists, including leaching of uncured oligomers and uncontrolled adsorption of small compounds. To alleviate these problems, we propose a new substrate for organ-on-chip devices: Polylactic Acid (PLA). PLA is a material derived from renewable resources, and compatible with high volume production technologies, such as microinjection moulding. PLA can be formed into sheets and prototyped into desired devices in the research lab. In this article we uncover the suitability of Polylactic acid as a substrate material for Microfluidic cell culture and Organ-on-a-chip applications. Surface properties, biocompatibility, small molecule adsorption and optical properties of PLA are investigated and compared with PDMS and other reference polymers.

**Significance:** Organ-On-Chip (OOC) technology is a powerful and emerging tool that allows the culture of cells constituting an organ and enables scientists, researchers and clinicians to conduct more physiologically relevant experiments without using expensive animal models. Since the emergence of the first OOC devices 10 years ago, the translation from research to market has happened relatively fast. To date, at least 28 companies are proposing body and tissue on-a chip devices. The material of choice in most commercial organ-on-chip platforms is an elastomer, Polydymethyloxane (PDMS), commonly used in microfluidic R&D. PDMS is however subject to poor reproducibility, and absorbs small molecule compounds unless treated. In this study we show that PLA overcomes all the drawbacks related to PDMS: PLA can be prototyped in less than 45 minutes from design to test, is transparent, not autofluorescent, and biocompatible. PLA-based microfluidic platforms have the potential to transform the OOC industry as well as to provide a sustainable alternative for future Lab-On-Chip and point-of-care devices.

## Introduction

Organ-On-Chip (OOC) is the convergence of two emerging research areas, microfluidics and tissue engineering, meeting the requirements for more physiologically accurate human tissues. OOC devices provide advanced models for cell and tissue culture, accelerating and facilitating the understanding of human biology, toxicology, disease progression and prognosis, while overcoming important ethical, financial and interspecies variances related to animal experimentations [1].

Since the birth of the first OOC device in 2007 [2], complex functionalities have been added to devices, leading to the so-called body-on-a-chip or human–on-a-chip fields. The translation from the research to the market was relatively fast. To date, at least 28 companies are proposing body and tissue on-a chip devices [3]. The material of choice in most organ-on-chip platforms has been poly(dimethylsiloxane) (PDMS), a thermo-curable elastomer. The majority of companies present in the OOC market (57%) sells PDMS products (Suppl. Inf. 1). Although PDMS is biocompatible, transparent, gas permeable, flexible, and relatively easy to manufacture at small scale, some issues have been encountered by consumers while using PDMS devices for certain applications. These problems include channel deformation, high evaporation, leaching of uncured oligomers, absorption of hydrophobic compounds and unstable surface treatment which can lead to inconsistent and unpredictable results with respect to some biological outcomes [4]. While some of these issues have been overcome (e.g. via soxhlet extraction in ethanol, or other organic solvent, to remove uncrosslinked oligomers [5]), PDMS moulding still remains a difficult process to fully automate [6] and significantly slows down the translation from the research to the mass market production.

To alleviate the drawbacks of PDMS, recent studies have focused on the use of thermoplastic as well as 3D printable materials. Poly methyl methacrylate (PMMA), Polycarbonate (PC), Cyclic olefin polymers and copolymers (COP and COC respectively) and Polystyrene (PS) are some common materials proposed from different suppliers as scalable alternatives. These thermoplastic materials present the advantage of being relatively cheap, and could translate easily to the mass market. However, these materials do not always exhibit good biocompatibility and are fossil-based, therefore unsustainable.

In a recent study, we proposed Polylactic acid (PLA), a biocompatible thermoplastic material from renewable resources and widely applied to tissue engineering, as new substrate material for the production of environmentally sustainable, single-use microfluidic devices. We optimised PLA workability in conjunction with a layer-by-layer laser-based prototyping technique and demonstrated basic microfluidic functions [7]. Here we provide evidence that PLA is a suitable replacement to other polymers in Organ-on-a-chip applications. We investigate surface properties, biocompatibility, absorption and adsorption of small compounds (< 900 Da), and optical properties of PLA in comparison to PDMS and with other reference polymers such as polystyrene. Finally, we provide an example of cell-tracking through a PLA-based organ-on-chip device.

## Results

### PLA functionalisation

An OOC device provides support for tissue attachment and organization to simulate organ-level physiology. The first interaction between the cells and the device happens at the material surface. The requirements for a suitable substrate material for healthy cell environment and tissue adhesion, are wettability, surface roughness and chemical composition[8], [9]. PLA is a hydrophobic polymer, which is a limiting factor for cell attachment. To address this, surface treatments are required to increase the substrate wettability [10], [11]. The most widely utilised and preferred surface modification methods rely on oxygen plasma and UV-ozone treatments [12]. However, several studies have reported the non-permanent nature of those functionalization methods, leading to the short term recovery of the hydrophobic property [13]. Such a phenomenon is particularly exhibited by PDMS, with a drastic increase of the contact angle one week after the treatment [13]. To overcome the hydrophobic recovery issue, PDMS devices need to be used just straight after the functionalisation process, which further limits their industrial applications. The contact angle value gives a quantitatively measure of the hydrophilic or hydrophobic nature of a surface. In particular, a material is hydrophilic if it has a contact angle below 90°, and hydrophobic if the contact angle is above 90°. Our PLA sheet manufacturing protocol (compression moulding protocol available in [7]) enables to produce slightly hydrophilic surface with a weak wettability. The measured contact angle is about 77-80°(Fig. 1A), while the desired contact angle for cell culture applications is 45°±4° (appropriate contact angle to avoid electrostatic interactions with hydrophilic molecules [13]). In order to permanently modify the surface properties of pristine PLA (*p*PLA), a wet-chemistry approach was adopted. A Sodium Hydroxide (NaOH) solution was employed to achieve an alkaline surface hydrolysis allowing the ester linkages of the PLA chains at the surface to produce carboxylic (-COOH) and hydroxyl (-OH) functional groups (Fig.1A). Using this functionalisation process, the final contact angle can be tuned by changing the functionalisation time and the solution concentration (Fig. 1B). This functionalisation protocol is stable over time, with no evidence of a hydrophobic recovery effect, even 9 months after the functionalization, at room temperature storage (Fig. 1C.i). In accordance with previous results reported by Tham et al. [14], this method acts on hydrolysing and eroding the material surface, inducing a visible change in the surface roughness (Fig. 1C.ii-D-E). In conclusion, the proposed functionalisation protocol allows one to tune the surface wettability of PLA to obtain the desired contact angle for cell culture. The resulting formation of the -COOH and of -OH groups at the surface can be readily used for further conjugation of surface modifying species in order to provide the best environment for the specific cell culture application [11], [14].

**Figure 1.**
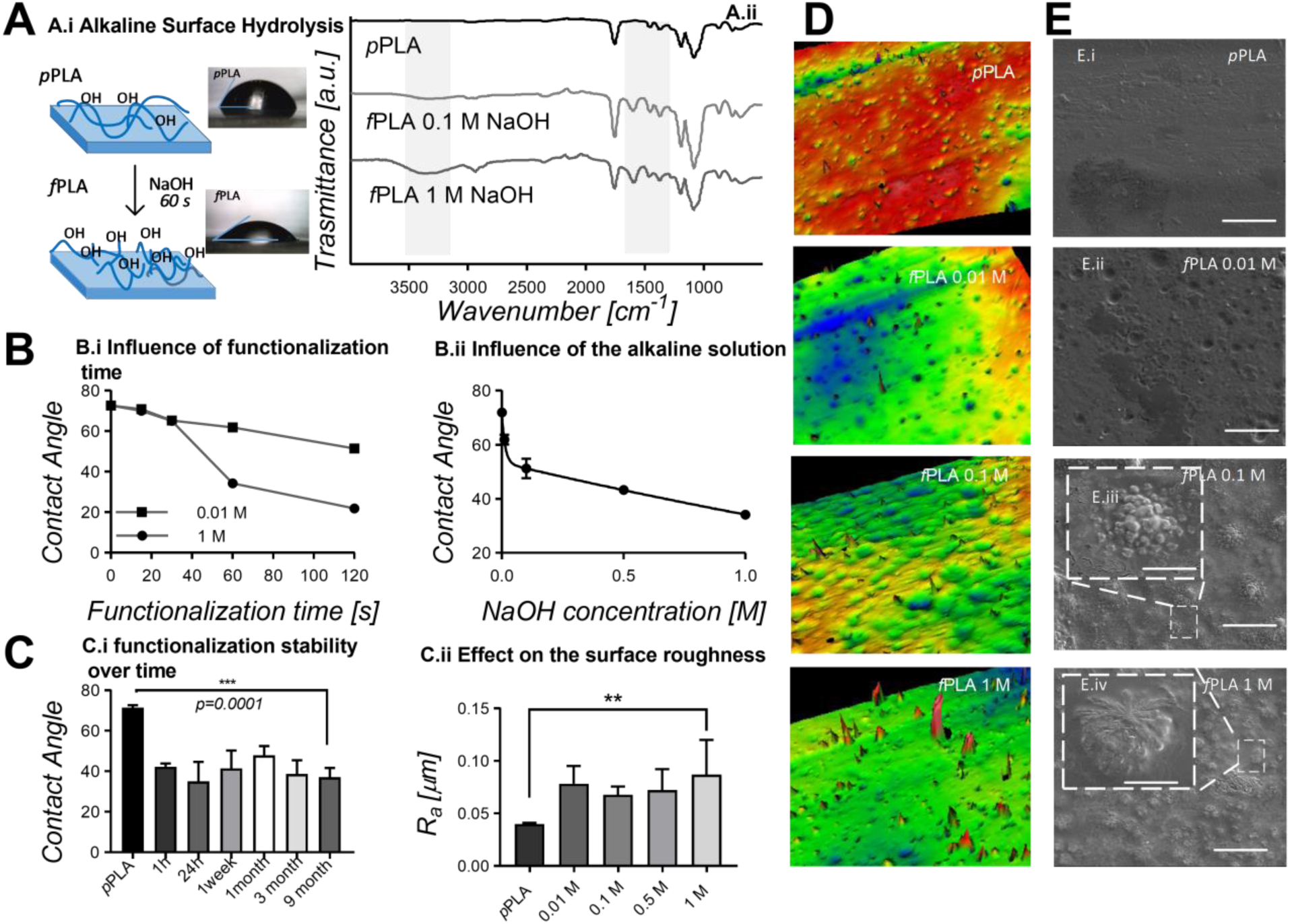
Tuning the surface properties of PLA. A.i) Schematization of the functionalization mechanism via Alkaline Surface Hydrolysis with photographs of a DI water drop on the pristine PLA substrate (top) and on the functionalized PLA substrate (bottom) A.ii) FTIR spectrum of pristine PLA (black curve), functionalized PLA using a concentration of 0.1 M of NaOH solution (light grey curve), and a concentration of 1 M of NaOH (grey curve). The grey areas underline the areas where a change in the spectrum is observed due to the functionalization. The first one (from left to right) at 3200 cm^-1^ is indicative of the formation of hydroxyl groups, the second one highlights a stretching of the C-C groups indicative of a partial hydrolysis of the treated surface. B.i) Influence of the functionalization time on the final contact angle for two different solution concentration, 0.01 M (box) and 1 M (circle). The two connecting lines are just to guide the eye. For all the points the error bars are shorter than the height of the symbol. B.ii) influence of the NaOH concentration on the final contact angle after 60 s of functionalization. The experimental results are fitted with a two-phase exponential decay function. For some points the error bar is shorter than the high of the symbol. C.i) Stability of the functionalization method adopted over nine months. C.ii) Effect of the solution concentration on the final surface roughness of the functionalized substrate after 60 s functionalization. D) Interferometer pictures of the unfunctionalized and funcionalized PLA surfaces with different solution concentration after 60 s functionalization time. Interestingly, a pillar formation structure is noticed with an increase of the pillar density with respect to the solution concentration. While increase of surface roughness is visible on SEM imaging, the pillar formation is partly damaged by the conductive gold layer. E) SEM pictures of the unfunctionalized and functionalized PLA surfaces with different solution concentration after 60s functionalization time. Atomic Force Microscope Imaging of Pristine and Functionalized PLA was also used to confirm the pillar formation. Data shown in Figure SI1.

### PLA biocompatibility

Biocompatibility in OOC devices can be defined as the absence of toxic effects to the cells or tissue interacting with the material, as well evidence of optimal growth of the cultured cells or tissues. PLA has been widely used in medicine as material for resorbable sutures, orthopaedic implants, scaffold for tissue engineering and drug delivery systems [15]. Therefore, its biocompatibility has been well established by a number of studies, typically demonstrating lack of low inflammation after implantation and compatibility with surrounding tissue [16], [17]. In this study, we specifically aimed at comparing the biocompatibility of pristine PLA (*p*PLA) versus functionalised PLA (*f*PLA), as well as against other common organ-on-chip and microfluidic materials, namely, PS (control material), PDMS, PMMA, and PC. Cell proliferation and cell death of two cell lines, human hepatocellular carcinoma cells (C3A) and adenocarcinomic human alveolar basal epithelial cells (A549) were examined as per material and method session. At 48 hrs of cell culture of the C3A cells, only the substrates made of *p*PLA, *f*PLA and *f*PDMS performed comparably with the control (albeit the differences with other polymers were small), and at 7 day time point *p*PLA and *f*PLA showed a decrease in cell death of 6% and 10% respectively, with respect to the control (Fig. 2 A). For the A549 cells at 48 hrs of culture almost all the substrates performed comparably well to PS control with no significant statistical difference in the cell death, while at 7 days all polymers caused the same percentage of cell death as compared to control with the exception of PLA that demonstrated a decrease in cell death (Fig 2 B). Cell proliferation data is shown in Figure SI2.

**Figure 2.**
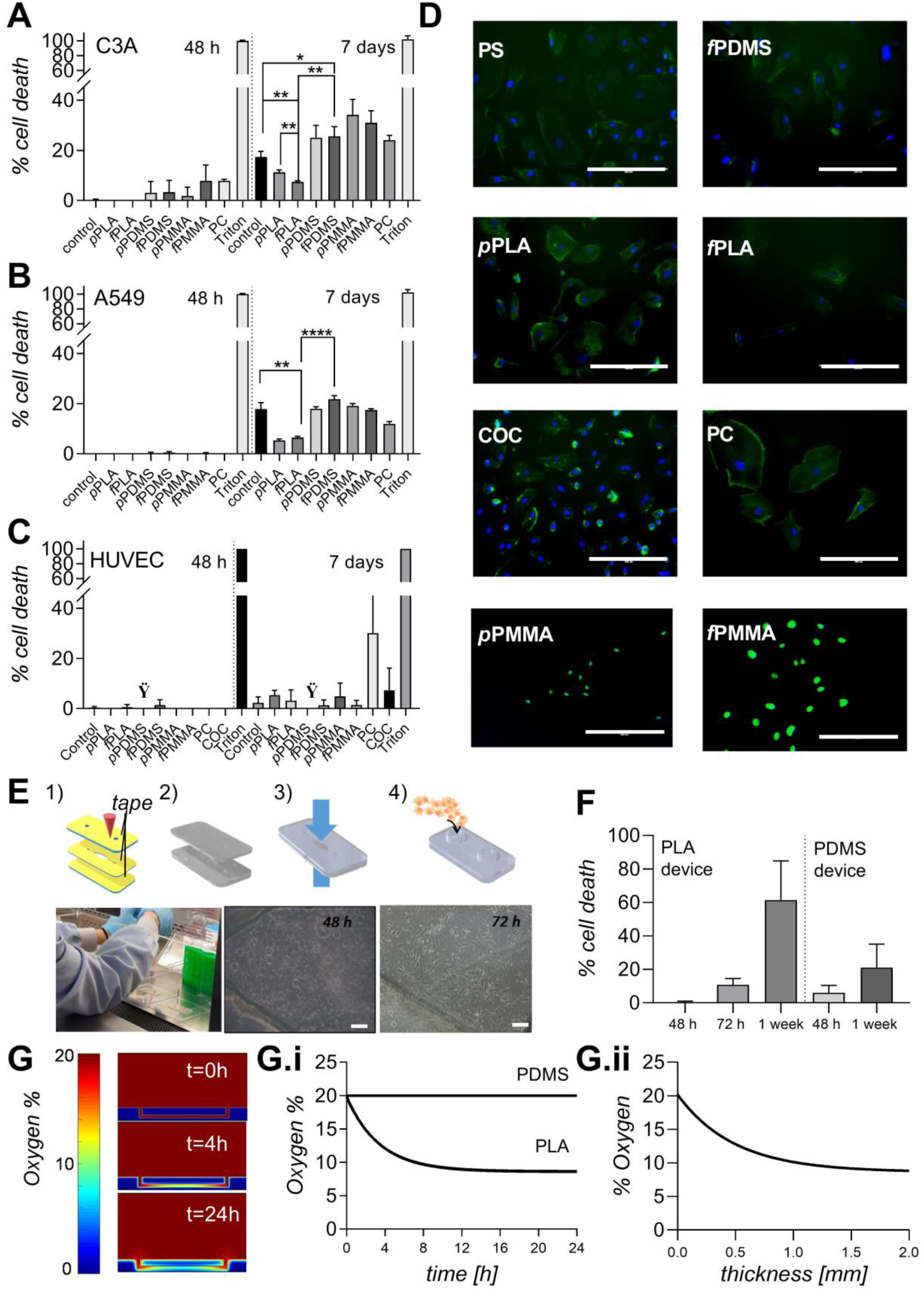
Assessing the material biocompatibility. A) Percentage of cell death of human hepatocellular carcinoma cells (C3A) on the different substrates at 48h and 7 days time points; B) Percentage of cell death of adenocarcinomic human alveolar basal epithelial cells (A549) on the different substrates at 48h and 7 days time points. C) Percentage of cell death of HUVECs cells on the different substrates at 7 days time point. The cells did not adhere on the pristine PDMS (Noted with a symbol: Ÿ). D) DAPI-actin staining of VEC cells at 7 days time point on the different substrates (scalebar is 200 µm). The transparency of PLA substrate enable imaging comparable to the rest of the other materials. E) Schematization of the protocol to manufacture the microfluidic device for culturing HUVECs, 1) CO_2_ laser cutting the 3 using the SLAM approach; 2) Cleaning and functionalizing the layers; 3) UV bonding the layers, 45 s excitation time and then holding the part in place at 55 C for 5 minutes; 4) Device ready to be sterilized and used. Underneath, from right to left, the photographs are showing the cells being loaded inside the device and bright-field microscope photograph of healthy HUVECs cells with good confluence after 48 and 72h culture, scale bar is 200 µm. F) Comparison of HUVEC cell death in the PLA and in the PDMS device. G) 2Dimensional simulation of the oxygen levels inside a PLA device at 0, 4 and 24 h culture time. The scale from 0 to 20% represent sthe oxygen level in the device and environment G.i) Simulated oxygen levels in function of culture time inside the PDMS and PLA microfluidic device. The oxygen level in the PLA level quickly reduces to 10% after a few hours G.ii) Simulated oxygen level in function of different thicknesses of PLA device sealing layer at 24h. A thickness of 0.250 mm would enable oxygen levels to be within 5% of the atmospheric level after 24h.

We also independently investigated the PLA biocompatibility with primary cells, which are sensitive cells with a finite life span and a limited expansion capacity, compared to cancer cell lines. We cultured freshly isolated human umbilical vein endothelial cells (HUVECs) for 7 days on the same substrates, with the addition of COC. Both *p*PLA and *f*PLA performed as the control PS (no statistical difference). PC and COC showed the higher cell death percentage (Fig. 2 C,D). Overall, the toxicity data of cancer and primary cells, indicates that PLA is a suitable material for cell culture. Translating from a polystyrene substrate to a PLA substrate was found to be marginally better for A549 and C3A cells, and similar from HUVEC cells.

To further evaluate the biocompatibility of *f*PLA and its suitability as substrate material for microfluidic cell culture and OOC application, a 3 layer microfluidic device consisting of a single cell culture rectangular chamber was fabricated as per material and method session (Fig. 2 E). Primary HUVECs were cultured for one week to evaluate morphology and proliferation. The transparency of the device allowed cell visualization (as shown in Fig. 2 E) and the cells easily adhered onto the PLA microfluidic walls without the need for adhesive protein coating. They formed a confluent layer after 3 days in culture, maintaining their natural morphology (Fig. 2 E). The material transparency and absence of autofluorescence enabled to easily evaluate cell viability using the ReadyProbes® kit. We compared the performance of the PLA device with an identical single chamber PDMS device. No statistically significant differences were noticed in the culture of the cells between the two different substrate materials, up to 72 h. At 1 week time point a significant increase in cell death was noticed inside the PLA device. This is likely to be related to the lower oxygen permeability of PLA with respect to PDMS, static condition of the culture (media changed every 24h) and the increasing oxygen demand due to cell proliferation [18].

In order to evaluate the oxygen concentration inside the PLA cell culture chamber and compare it to a PDMS chamber, we used FEATool Multiphysics, a Matlab toolbox to create a 3D model. A time dependent condition was imposed and the oxygen concentration simulated over 24h with a time step of 1h assuming an oxygen consumption rate of the cells of 0.37×10^−4^ mol/s [19]. We carried out different simulations by changing the thickness of the top sealing layer (Fig. 2G) and plotted the oxygen concentration level at 24 h time point inside the microfluidic chamber in function of the thickness of the layer. The results from the numerical simulation suggests that using a 0.1 mm top sealing layer allows an oxygen level at 24h comparable to a PDMS device with a 2 mm thickness. Alternatively, continuous perfusion of media may be used. Further studies taking oxygen permeability, and design variable into account will be undertaken.

### Absorption and Adsorption of small molecules on PLA substrates

Several reports have shown that, unless treated, PDMS can absorb small hydrophobic compounds such as steroids and hormones [20]–[24]. This behaviour can be explained by the porous and hydrophobic nature of its polymeric network. This phenomenon can be a major drawback in bioassays, in particular drug testing assays where drugs engineered for rapid delivery are typically smaller than 500 Da and diffuse uncontrollably inside the microfluidic device [21], [22]. Several methodologies have been proposed to overcome this issue; however, they result in delays in prototyping or manufacturing time. On the other hand, PLA, as a thermoplastic material should not show any absorption behaviour, but it can be subjected to surface adsorption. In this study, the absorption of both hydrophilic and hydrophobic compounds was assessed in laser-cut *p*PLA and *f*PLA and PDMS channels. To test the absorption of hydrophobic compounds, Nile Red, a small hydrophobic fluorophore (318.37 Da), was selected as a representative molecule, while fluorescein salt (376.27 Da) was selected to test the absorption of a typical hydrophilic compound. The devices have been designed and manufactured with a channel cross-section of 0.8 mm^2^ (Fig. 3A). A fluorescence image of each empty channel was taken with a microscope in order to create the null absorption reference point (Fig. 3B). To create the maximum absorption reference point, the channel was filled with the fluorescent solution (Fig. 3B). The solution was incubated inside the channel for 60s and then aspirated out of the channel from the outlet. Each channel was then cleaned with deionised water if loaded with Nile Red or with ethanol if loaded with fluorescein salt solution. A picture was taken after 60s of the cleaning channel cycle and the fluorescence intensity measured with the imaging software ImageJ and normalised with respect to the reference points. The rinsing-cleaning-imaging cycle was repeated 20 times (Fig. 3B) and the normalised fluorescence intensity was plotted against the number of cycles (Fig. 3C). This experiment showed PLA does not absorb hydrophobic or hydrophilic compounds, even when functionalised, while PDMS predictably absorbs small hydrophobic compounds (reaching 50% of absorption after 20 cycles), making PLA a superior material in this respect.

**Figure 3.**
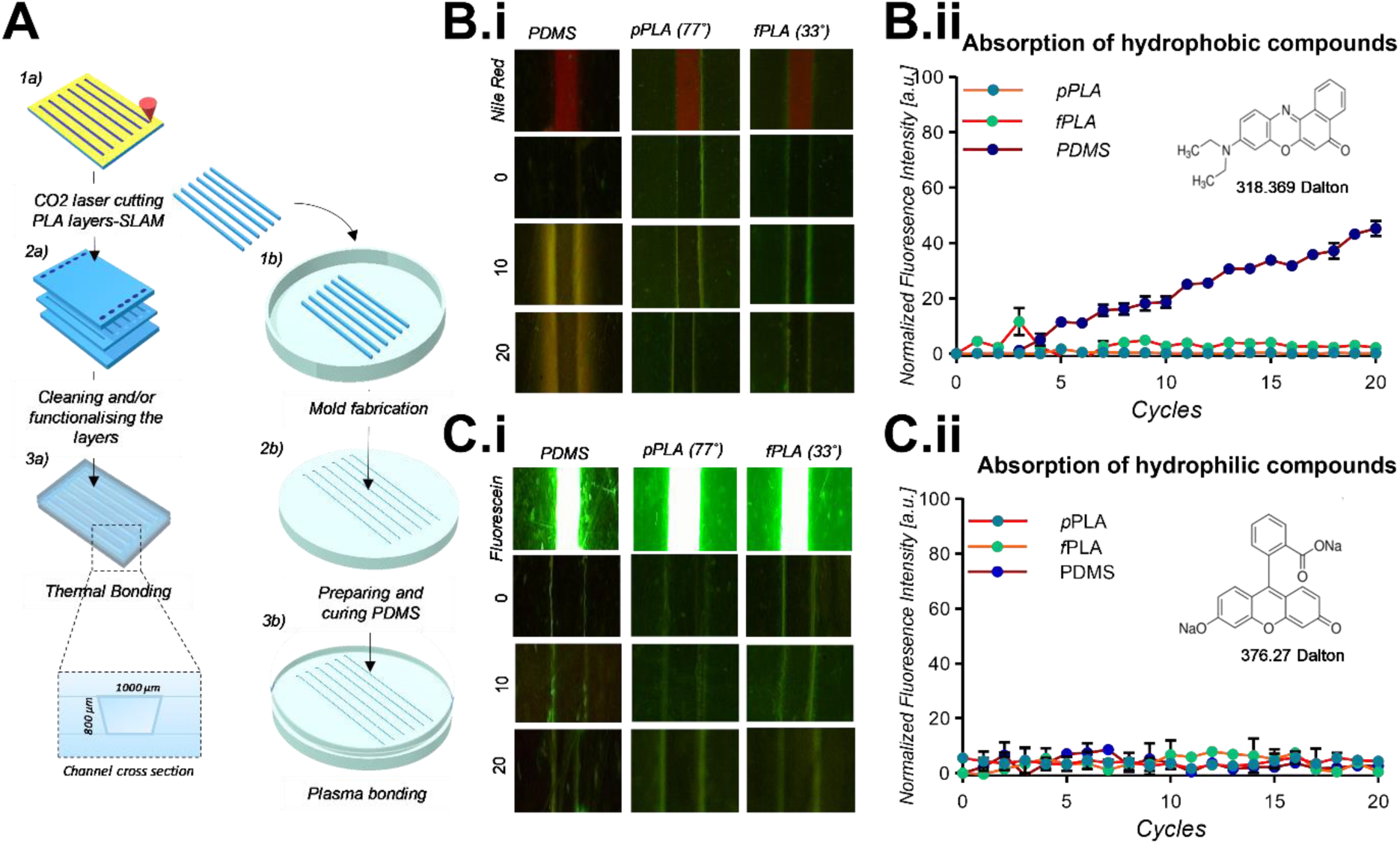
Investigation into the adsorption of small molecules in PLA substrates. A) Schematization of the protocol to manufacture PLA and PDMS microdevices. 1a) SLAM method is used to cut a channel of 1 mm width and access holes from a 1mm PLA sheet for channel and top layers, respectively. 2a) Top, bottom and channel layers are cleaned and functionalised for 60 s with 1M NaOH solution.3a) The layers are bonded together at 70°C for 1h using binder clips to maintain the layer alignment 3a). In insert the channel cross section. 1b) The cut PLA parallelepipeds from 1a) are used as negative mould to fabricate the PDMS device to have the same channel cross-section for the different devices. 2b) The PDMS is mixed and placed in the mould and cured. 3b) Plasma bonding of the PDMS device. B) Adsorption of hydrophobic compounds. B.i) top view of the channel for (from left to right) PDMS, pristine PLA and functionalized PLA. From the top to the bottom picture of the channel filled with a solution of Nile Red in ethanol (1µM), representing one of the reference point; then picture of the empty channel, representing the 0 reference point, and picture of the channel at the 10^th^ and 20^th^ cycle. B.ii) Normalized fluorescence intensity, measured at the centre of the channel, plotted in function of the cycle number. In the insert molecular structure and molecular weight of the Nile Red. C) Absorption of hydrophilic compounds. C.i) Top view of the channel for (from left to right) PDMS, pristine PLA and functionalized PLA. From the top to the bottom picture of the channel filled with a solution of fluorescein salt in DI water (1µM), representing one of the reference point; then picture of the empty channel, representing the reference point, and picture of the channel at the 10^th^ and 20^th^ cycle. B.ii) Normalized fluorescence intensity, measured at the centre of the channel, plotted in function of the cycle number. In insert molecular structure and molecular weight of the fluorescein salt.

### Optical properties of PLA: Transparency and autofluorescence

In order to study the organ functionality and biological responses at the cellular and molecular level, the optical properties of an organ-on-chip system substrate material, are of fundamental importance and relevance [25]. It is well established that PDMS, PMMA, PC, COP and COC have good or excellent optical properties and transparency. Furthermore, they have a low fluorescence background or autofluorescence, when irradiated near UV wavelength, a desirable feature for fluorescent imaging [26]. The optical properties of PLA have been investigated and reported previously [27], however, very little is known about PLA autofluorescence properties. Here the transparency and autofluorescence of PLA substrate was investigated, analysing possible changes due to functionalisation (*p*PLA versus *f*PLA) in comparison with PDMS, PMMA, PC and COC substrates. We observed no difference between *p*PLA and *f*PLA with regards to transparency as transmittance of the light in the visible region (Fig. 4A). *f*PLA showed 92% of transparency, higher than PDMS, but lower with respect to the other materials tested. While PMMA showed the highest transparency (96 %), followed by COC, PC, PLA and PDMS (Fig. 4 B).

**Figure 4.**
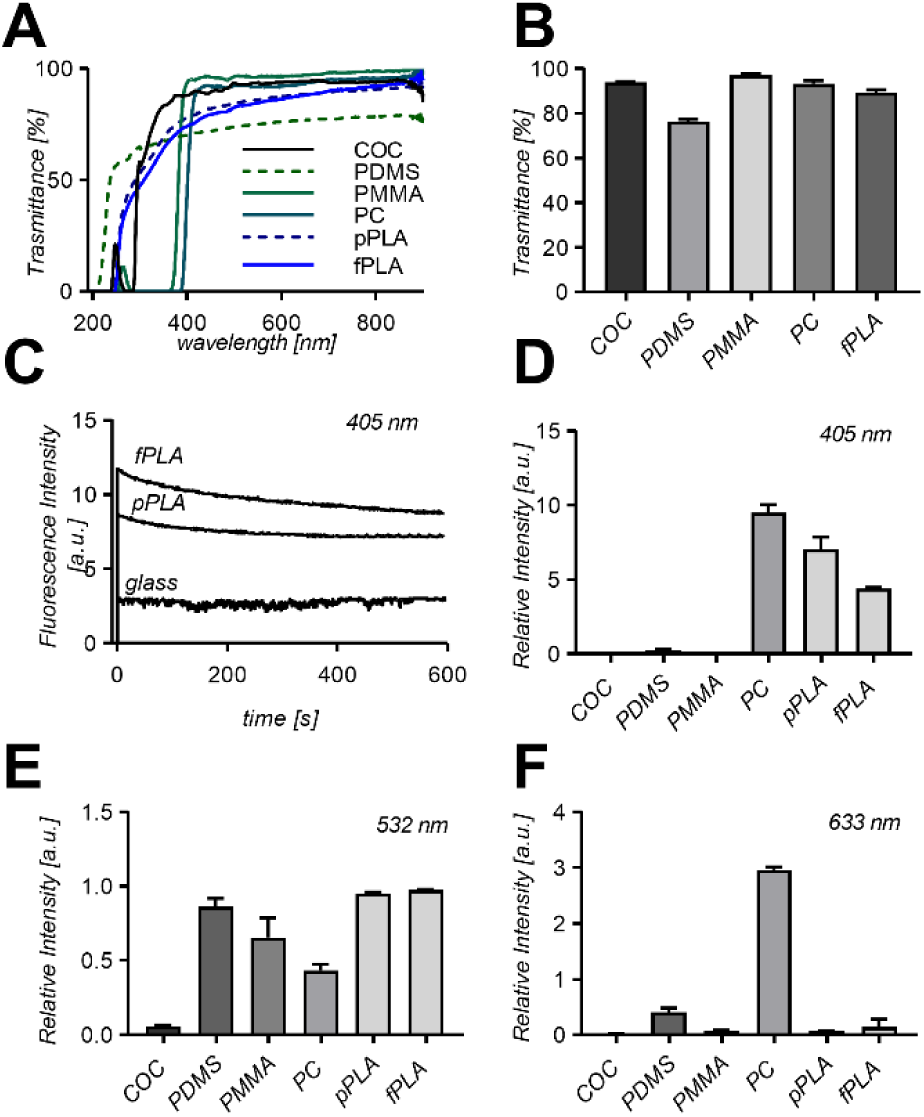
Investigation into PLA optical properties: A) UV-VIS spectrum of PLA (functionalised: fPLA and pristine PLA: pPLA) and different COC, PDMS, PMMA and PC materials; B) Comparison of light transmittance in the visible region of fPLA compared to COC, PDMS, PMMA and PC substrate materials. C) Fluorescence decay profile under 600s continuum illumination of functionalised PLA, pristine PLA and glass; Autofluorescence intensity relative to glass of the different substrate materials after 60 s continuum illumination with 405 nm excitation D), 532 nm excitation E) and 633 nm excitation F).

In OOC applications fluorescence imaging is often required over a period of time rather than instantly [28]. Hence, the analysis of the autofluorescence over time under illumination is highly beneficial. Therefore, next, the autofluorescence was measured over a period of 600s (Fig. 4C). *p*PLA and *f*PLA were analysed and compared to glass, with data showing a gradual fluorescence decay for both. Contrary to the transparency behaviour, a notable difference was noted between the pristine and the functionalised PLA, with *f*PLA showing a higher autofluorescence due to the increased presence of –OH groups in the functionalised sample. The use of material without defect such as bubbles or lamellae is of importance to the reduction of the background autofluorescence. Some examples of defects can be found in Fig SI3. Furthermore, the autofluorescence of *p*PLA and *f*PLA were measured and compared to COC, PC, PDMS and PMMA with reference to glass. In these experiments, different wavelengths were tested, covering the range used in fluorescence microscopy. The fluorescence background was taken after 60s illumination [26]. As expected, for all the materials tested, the autofluorescence decreased with increasing wavelengths. An exception to this was PC, which demonstrated a lower fluorescence background at 532 nm. PLA showed autofluorescence levels no higher than ∼ 1 times glass levels and comparable to the other tested materials (Fig. 4 D,E,F). Although the numerical fluorescence background measured for both pristine and functionalised PLA were higher than other materials, these results are not hindering PLA suitability for fluorescence imaging [26], [28].

### Cell tracking in a PLA Organ-On-Chip device

The role of an Organ-On-Chip device is not only to culture different cell types successfully but also to provide a micro-environment for cells, typically superior to standard cell culture in wells. To this end, Organ-On-Chip devices can be engineered in order to guide and spatially confine the cells. Recently, Biselli et al.[25] used an OOC device as powerful tool to study and quantitatively analyse the cancer-immune cells interactions via imaging analysis. The velocities, turning angle and path of the moving cells were used as indicator to study anticancer immune responses. Having demonstrated that the transparency of *f*PLA substrates is comparable to that of other common polymers used in microfluidic, we designed and manufactured an Organ-On-Chip device for cell tracking in a confined channel. A device composed of three layers was manufactured using a combination of hot embossing and Sacrificial Layer laser Assisted Method (SLAM) approaches from *f*PLA substrates [7] (Fig. 5 A). In this experiment, prostate adenocarcinoma PC-3 cells were cultured on the device and a time lapse video was recorded using a customized inverted microscope (as per material and method session) in a CO_2_ incubator at 37°C. A microphotograph of the region of interest was taken at 1 minute intervals using a 4x magnification objective and videos analysed with a custom written software as per material and method session. Kinematic descriptors such as speed, mean curvature, and angular speed were extracted and compared for a PS Petri dish, PLA disc or inside a device. The kinematic descriptors between the PLA and the PS substrates were very similar despite slight difference of surface topology [29]. Additionally, the diffusion coefficient of individual cells was comparable on both substrates, thus demonstrating that cells on PS and PLA substrates have the same behaviour (Fig. 5B). On the other hand, the kinematic descriptors extracted from cell tracking in the PLA OOC device showed cell motility differences compared to the previous bare and open substrates (Fig. 5C). Cell random walk was observed, but as expected, subdiffusivity was noted in the confined environment. This might be made worse due to the difference in the oxygen permeability of the material. The differences observed in cell motility could be overcome by providing a better surface topology, a protein coating in culture channels to improve cell adhesion, and reducing the thickness of the device or applying media flow for gas exchange [30]. In conclusion, the transparency of *f*PLA substrates allowed advanced cell tracking in the device, similarly to the control PS Petri dish, and we were able to observe a random cell confined walk in PLA OOC device (Fig. 5D).

**Figure 5.**
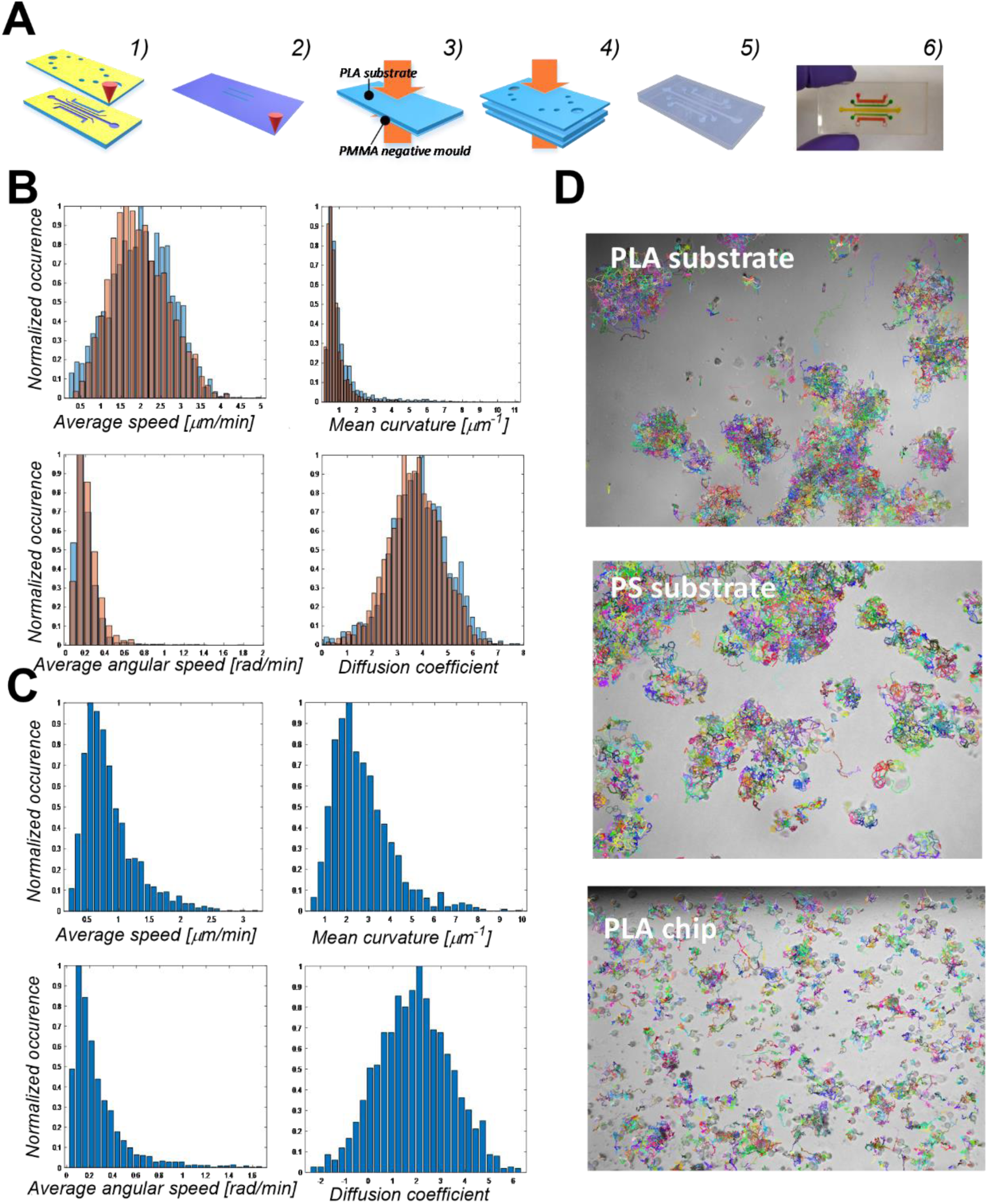
Demonstration of PLA suitability for cell tracking in Organ-on-a-chip and microfluidic cell culture devices. A) Schematization of the protocol to manufacture PLA Organ-on-a-chip or microfluidic devices. A-1) SLAM approach to create inlet and outlet holes in the top layer and to microstructure the channel on the middle layer. A-2) Rastering a PMMA substrate to create a mould for the bottom layer with diffusion channels. A-3) hot embossing the structure from the PMMA mould to PLA, using a temperature of 70 °C and a pressure of 0.5 MPa for 15 minutes. A-4) NaOH functionalization and UV activation of the surfaces with thermal bonding of the different layers. T=50 °C for 5 minutes. A-5) The assembled device is cooled down while maintaining the layers in contact. A-6) Photogragh of the final device filled with red, green and yellow food dyes B) Statistics of the kinematic descriptors for PC-3 prostate cancer cells on a PLA open disk (blue bars) and on a PS well plate (orange bars) B.i) Speed, B.ii) Mean curvature B.iii) Angular speed and B.iv) Diffusion coefficient. C) Statistics of the Kinematic descriptors of the PC-3 prostate cancer cells on a PLA Organ-On-Chip device C.i) Speed, C.ii) Mean curvature C.iii) Angular speed and C.iv) Diffusion coefficient. D) Extracted PC-3 prostate cancer cells trajectories through the Cell-hunter software on a disc of PLA D.i), on a PS substrate D.ii) and on a PLA device.

## Discussion

We introduced PLA as a novel material for Organ-On-Chip devices. PLA can be machined via laser-cutting as shown in this article, or by 3D printing as shown elsewhere. In this paper, we showed that PLA can offer many distinct advantages as a substrate for Organ-On-Chip applications over other polymers. First, PLA could be functionalised via a rapid (60s) functionalisation process using Sodium Hydroxide, reducing its native contact angle of about 70 to a contact angle of 40°, suitable for cell culture. We found this surface treatment to be stable over at least 9 months, which shows commercial PLA devices treated in such a way would have excellent shelf-life. Secondly, cells could be cultured on PLA surfaces, as easily as on conventional polystyrene (PS) petri-dishes. Pristine and functionalised PLA performed better with regards to cell proliferation and cell toxicity compared to the majority of uncoated polymeric material such as PMMA, PC, COC. Thirdly, we demonstrated that PLA performed significantly better than PDMS with regards to the adsorption and absorption of small molecules. Taking a hydrophilic representative molecule, Nile Red, of same size as oestrogen, a classic target compound in cell studies, we showed that a PDMS adsorbed 50% of the Nile Red after 20 load and wash cycles, while a similar PLA channel showed no adsorption, whether or not it had been functionalised. Finalised, we showed that lab-made PLA substrates are highly transparent and present very little auto-fluorescence, enabling high quality imaging, which is demonstrated through a cell-tracking experiment in an Organ-On-Chip device. Finally, advanced biological protocols such as random walk cell tracking were demonstrated in a PLA device.

PLA is compatible with a range of complex integration, such as conducting electrodes, which we demonstrated in a previous publication [7]. While here we used a layer-by-layer approach to prototype Organ-On-Chip devices, PLA can be also formed, moulded, and printed. PLA is a popular material for filament deposition modelling, or 3D-printing and 3D printed microfluidic devices have demonstrated elsewhere [31]. However, these PLA 3D-printed devices currently lack the transparency necessary to enable high-resolution imaging and additional features are required to circumvent the lack of transparency. Here we showed that high PLA transparency can be achieved using moulding and laser-cutting techniques, which applies to both prototyping and high-volume manufacturing.

Up to now, other than ease of prototyping, one of the main reason for choosing PDMS for OOC application was its good oxygen permeability, with a diffusion coefficient of 3.5*10^−9^ m^2^/s. In our cell culture experiments, we found the PLA oxygen diffusion coefficient of 2.5*10^−12^ m^2^/s to be a limiting factor. However, continuous media perfusion or thinner lid are expected to remove this limitation as shown by our simulation results. Additionally, PLA lower oxygen diffusion coefficient enables the reduction and better control of water evaporation, and limit issues arising from osmolality differences induced by PDMS high oxygen diffusion coefficient. In this article PLA sheets were produced in the lab and did not have a complete planar topology. It is anticipated that industrial PLA substrates will show improved surface topology and even better optical properties.

In summary, we have demonstrated the suitability of PLA as a new substrate material for Organ-On-Chip devices. Our approach is designed to address many disadvantages inherent to conventional PDMS microfabrication technique. In conjunction with its biocompatibility, inertness to small molecules, optical qualities, PLA microfluidics will open opportunities to develop more sustainable solutions for Organ-On-Chip and microfluidic laboratories, in research or in industry. PLA represents a more environmentally sustainable solution than other thermoplastic fossil-based materials proposed in the market as alternative to PDMS and this study offers a novel alternative solution for completely sustainable devices. Ultimately, we envision that microfluidic device engineers will embrace a Design for Sustainability approach, and look into biopolymers, such as PLA, to develop sustainable single-use devices, for a wide range of applications, from Organ-On-Chip to Point-Of-Care devices.

## Materials and methods

### Materials

Polylactic acid (Natureworks® 2003D) was purchased from Naturework in pellet format. PMMA Clarex® was purchased from Weatherall Ltd, in a 1 mm sheet form. PDMS Sylgard 184 was purchased from Dow Corning, while COC (Topas), PC (RS components) were acquired in 1 mm sheet. All the reagents used were purchased from Sigma Aldrich, unless specified in the text.

### PLA Functionalization

The functionalisation process was an alkaline surface hydrolysis. NaOH (sigma Aldrich) was dissolved in DI water to prepare 0.01, 0.1, 0.5, and 1 M solution concentrations. The Pristine PLA sample is dipped inside the solution and kept in for specified amount of time (0, 10, 30, 60, 120 sec). A contact angle of about 40°, recommended for cell culture application [8], [32], was achieved by dipping the substrate for 60 seconds. Then the sample was washed with water and dried with compressed air. To measure the effect of the functionalization time, and solution concentration, the contact angle of the samples was measured using a custom-made static contact angle apparatus [33] The images acquired via a Dinolite microscope (model) were analysed with ImageJ plugin contactj. Each measurement was carried out 6 times. To assess the chemical changes at the surface an attenuated total reflection, ATR-FTIR, analysis was carried out collecting 200 scans in the range of 4000-400 cm^-1^, with a resolution of 4 cm^-1^ using a Nicolet Is5 spectrophotometer. The effect of the surface functionalization on the surface were studied using a white light interferometer (Zygo) with 1 nm resolution in vertical direction. The Roughness (which one, Ra) quantitative determination was carried out using Metro.Pro 8.2. For morphology analysis SEM (Quanta FEG 650 SEM) images were acquired in high vacuum mode.

### PLA Biocompatibility

The human hepatoblastoma C3A cell line was obtained from the American Type Culture Collection (ATCC, USA). The cells were maintained in Minimum Essential Medium Eagle (Gibco, UK) with 10% foetal bovine serum (FBS) (Gibco, UK), 2 mM L-glutamine, 100 U/ml penicillin/ streptomycin, 1 mM sodium pyruvate and 1% nonessential amino acids (all Sigma, UK) (termed complete medium), at 37°C and 5% CO_2_. The A549 cells were purchased from ATCC and maintained in RPMI (GIbco, uk) with 10% FBS, 100 IU/ml non-essential AA, 2 mM L glutamine and 100 U/ml Penicillin/Streptomycin. For the biocompatibility experiments, both cell types were seeded in one 24-well plate at a cell density of 50,000 cells per well in 1 ml of complete medium. The medium was changed every 48 hrs up to a period of 7 days. Subsequent to appropriate cell culture period (48 hrs or 7 days) the cell supernatants were collected and frozen at −80°C and later used for the adenylate kinase assay. In the alamar blue assay (proliferation), the supernatant was removed from all well, and stored in a freezer -80°C as described above. 1 ml of diluted alamar Blue reagent (Sigma, UK) (0.1 mg/ml) reagent was added to all (Sigma, UK). The plates were incubated at 37°C for 90 minutes and fluorescence was measured at 544/590nm. The loss of cell membrane integrity was evaluated utilising a ToxiLight™ bioassay kit (Lonza, USA). Briefly, 20 µl of cell supernatant was transferred to a luminescence compatible plate before the addition of 80 µl of adenylate kinase (AK) detection buffer. The plates were incubated for 5 min at room temperature and luminescence quantified. These experiments included a positive control: polystyrene cell culture treated plastic and a negative control: 0.1% Triton X-100 (Sigma, UK).

### PLA device biocompatbility

Devices were sterilized using EtOH, filled with culture medium and equilibrated in incubator (37 °C, 5% CO_2_) for 3 h before cell loading. HUVECs P6 were cultured on flask in EGM™-2 Endothelial Cell Growth Medium-2 BulletKit ™ (Lonza). Cells were detached and loaded in PLA devices with seeding density 1 ×10^6^ cells/ml. Devices were placed in incubator (37 °C, 5% CO_2_) to allow cells to adhere to the surface for 30 min. Then, 200 μl medium was added to inlet and outlet ports, and replaced every 24 h. To evaluate cell viability, a live dead assay (ReadyProbes® Cell Viability Imaging Kit (Blue/Green)) was used after 3 days and 7 days of culture.

### Finite Element Analysis

Finite element analysis was conducted using FEATool Multiphysics™ version1.10 (Matlab) to determine the oxygen concentration level inside the microfluidic cell culture chamber. In the software, a simplified 2D model of the PLA microfluidic device was designed (75×3.6mm device, 25×0.8 mm cell culture chamber, with 2 mm inlet and outlet holes, 12×8mm reservoir placed on the top of the 2mm sealing layer). Furthermore, the convection diffusion equation was solved in the time-dependent domain. For the oxygen diffusion simulation, a constant flux threshold (20%) was applied on the external sidewalls of the device and of the reservoir. And an oxygen diffusion coefficient of 2.5×10^−12^, 3.5×10^−9^, 4×10^−9^ and 2.14×10^−5^ m^2^/s was used for PLA, PDMS, water and air respectively. Imposing the starting concentration level of Oxygen inside the material device of 0%, and of 20% in water. To simulate the oxygen consumption rate, a consumption of oxygen of 0.37×10^4^ mol/s was imposed on a rectangular region of (25×0.08mm) at the bottom part of the cell culture chamber.

### Organ-On-Chip device preparation

To manufacture the PLA Organ-On-Chip devices, firstly PLA sheets were manufactured using a manual hydraulic heated press as per method described in [7]. A prototyping approach was previously described [7]. Briefly, each layer was cut using a CO_2_ laser cutter (Epilog Mini Elix 30W) using SLAM approach, which consists of applying a thin tape (100 µm) on the material prior to laser-cutting. The use of a commercial CO_2_ laser enables the formation of channels with 200 µm minimum size, however, other types of laser enable smaller channel widths. Some examples are discussed in Figure SI4. Then the different layers are cleaned by sonicating in pure Ethanol. To manufacture the functionalized device, the cleaned layers are dipped in a 1M NaOH solution for 60 s, then cleaned with water, 2-Propanol (Sigma Aldrich) and compressed air. The surfaces are then chemically activated under UV exposure (254nm) for 45 s. The different layers are then bonded together placing them in a sandwich composed of two glass slides, and kept them in contact at 50 °C for 10 minutes. Once the assembly has cooled down to room temperature, it is disassembled and the device is ready to be tested.

### Small molecules absorption and adsorption

To assess the bio-inertness of the material, pristine PLA was compared with functionalised PLA and with PDMS. To manufacture the device, specifically 6 channels of 1 mm width and 0.8 mm depth were cut through the PLA layers. The remaining cut pieces were used as negative mould to manufacture the PDMS device. The PLA device were then assembled as per chip preparation section. To manufacture the PDMS device, PDMS was mixed with the curing agent with a weight ratio of 1:10 and poured into a PS petri dish where the negative mould had been previously attached using a double sided adhesive tape (3M tape 469 MP). The PDMS was then left for 1h30 at room temperature prior to being placed for 1h at 80°C on a heated plate. In the same way a PDMS layer with input and output access holes were prepared. The two layers were then bonded together using an oxygen plasma treatment for 2 minutes (Diener electronic plasma surface technology). To test the absorption of hydrophobic compound Nile Red (sigma Aldrich) was dissolved in EtOH 99% to obtain a solution concentration of 1 µM. To test the absorption of hydrophilic compounds fluorescein salt was dissolved in DI water (1uM). A fluorescent microscope (AM4115T-GFBW) was used to take the fluorescent images and the image analysis freeware Imagej was used to analyse the fluorescence intensity. An advantage of this absorption experimental set-up is that it is fast and it does not require specific apparatus.

### PLA Optical properties

The transparency of the materials was tested analysing the transmittance of the light in the visible region using a UV-VIS spectrophotometer (Shimadzu UV-2550). The autofluorescence of the plastic materials of interest was measured using a Leica SP8 3X STED laser scanning microscope equipped with two Internal Spectral Detector Channels (PMT). Current from the PMT was amplified and data was acquired using the software LAS X from Leica. Laser Kit WLL2 (supercontinuum white light laser [pulsed]), 200 excitation lines from 470-670 nm was used as light source. The fluorescence was filtered by suitable laser filters for 405 nm (emission filter 460/510 nm), 532 nm (emission filter 580/640 nm) and 633 nm (emission filter 650/720 nm). Materials were wiped with ethanol preceding the imaging and were attached to a microscope slide before placing them on the microscope stage. The microscope objective (20x) was focused at the bottom surface of the samples and the objective was moved towards them. Auto-fluorescence imaging was done after 60 seconds illumination of each sample at each laser wavelength of interest (405, 532 and 633 nm excitation wavelengths). For each sample three unbleached sample spots were identified and selected for illumination. The analysis of the material auto-fluorescence long term recovery was done by illuminating and recording the samples for 10 min at both 405 nm (emission filter 460/510 nm) and 633 nm (emission filter 650/720 nm) wavelengths

### Cell imaging and tracking on a PLA device

#### Cell Culture

PC-3 prostate cancer cells were cultured at 37°C in a 5% CO_2_ humidified atmosphere in RMPI (10% fetal bovine serum, 100,000 units/litre penicillin, 50 mg/L streptomycin and 200mM glutamine). Cell seeding in PLA support (1st method). PC-3 cells were seeded on PLA lab-on-chip at a final concentration of 20.000 cells/80µL. Every hour new RPMI fresh medium was slowly added to prevent the channels from drying out completely. Cell seeding in PLA support (2nd method). PC-3 cells were seeded on PLA lab-on-chip at a final concentration of 20.000 cells/80µL. Tips containing 20µl of RMPI were positioned at the end of each microfluidic channel to allow replacing the medium every 24 hours (instead of 1 hour).

#### Cell imaging

A customized microscope has been designed for the scope: the prototype consists of a small-scale inverted microscope suitable to work in high humidity environments (incubators) with factory standard optics, custom aluminium structure and fully sealed electronics. A custom firmware was implemented in Matlab2017a® environment in order to have full control on acquisition methods and light exposure. The microscope time setting used was one frame per minute for a total of 12hr. The captured images have a FOV of 1.2mm width by 1mm height with a spatial resolution of 0.33 µm/px.

#### Cell tracking

The video sequence is then processed by a proprietary software, Cell-Hunter elsewhere validated in cancer immune interaction [25], [34] and block replication in prostate cancer cells [35]. With the aim to speed up the cell localization process, images were downsampled at a spatial resolution of 0.66 µm/px. Typical dimension of cells involved allows us to avoid localization errors. The localization step is implemented by the Circular Hough Transform (CHT) method, under the assumption of circular-shaped objects, that simultaneously allows one to locate cell nuclei in a given image providing an accurate estimate of individual cell radii. Cell occlusion and overlapping problems are solved through the geometrical mechanism behind. After cell nuclei are located, an improved proximity tracking approach based on the Munkres’ solution to the non-square Optimal Assignment (OA) problem is applied. Track refining procedure related to the reliable assumption of low expected cell motility with respect to the very high frame rate achieved by the microscope allows to reduce false tracks and join small tracks in a whole.

#### Cell track kinematic characterization

For the sake of comparing the kinematic characteristics of cells involved in the competing experiments, we extracted standard descriptors such as average speed, curvature, and angular speed, along with the coefficient of diffusion [36]. The distribution of such descriptors computed over all the tracks extracted are shown in Fig. 5B and C.

## Acknowledgements

A.O is funded by a James Watt scholarship. AO, VP, EM and MKK acknowledge the Organ-On-A-Chip Technologies Network Sabbatical funding scheme. M.K.K. acknowledge funding from the Engineering and Physical Sciences Research Council, EP/R00398X/1. V.P. acknowledges the PROM project (Project number 748903), funded by H2020-MSCA-IF-2016 and the MRC Confidence in Confidence scheme. We thank Rory Duncan for access to the Edinburgh Super-Resolution Imaging Consortium (ESRIC) Facility.

## Author contributions

Author contributions: AO, NH, VLC, VP and MKK, designed the research; AO, AMC, DV, AK, DDG, AM, LG, KW, VM, VP performed research; VS and DH contributed to reagents and analytic tools; AO, AK, DDG, NH, EM, VP, MKK analysed data; and AO and MKK wrote the paper. All authors read and reviewed the paper.

The authors declare no conflict of interest.

## Supplementary Information

**Table S1.**
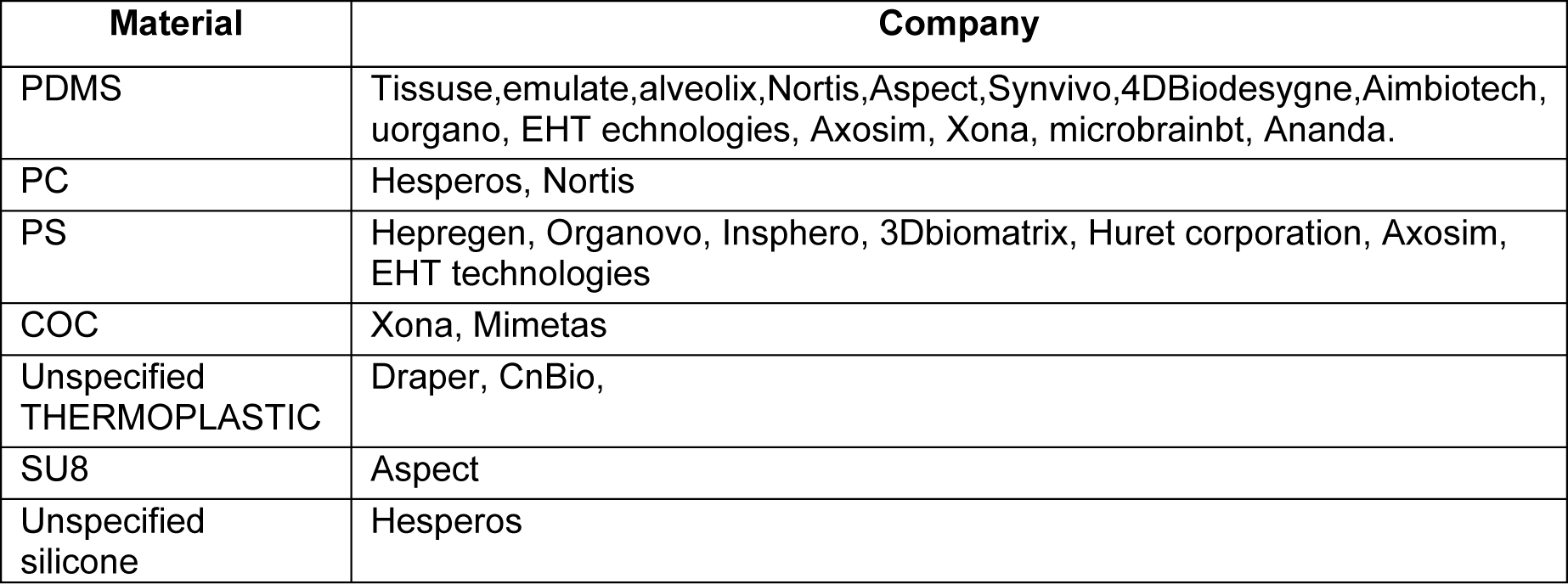
Materials used in the mass production of Organ-on-a-chip devices. The companies have been taken from [3]

**Figure SI1.**
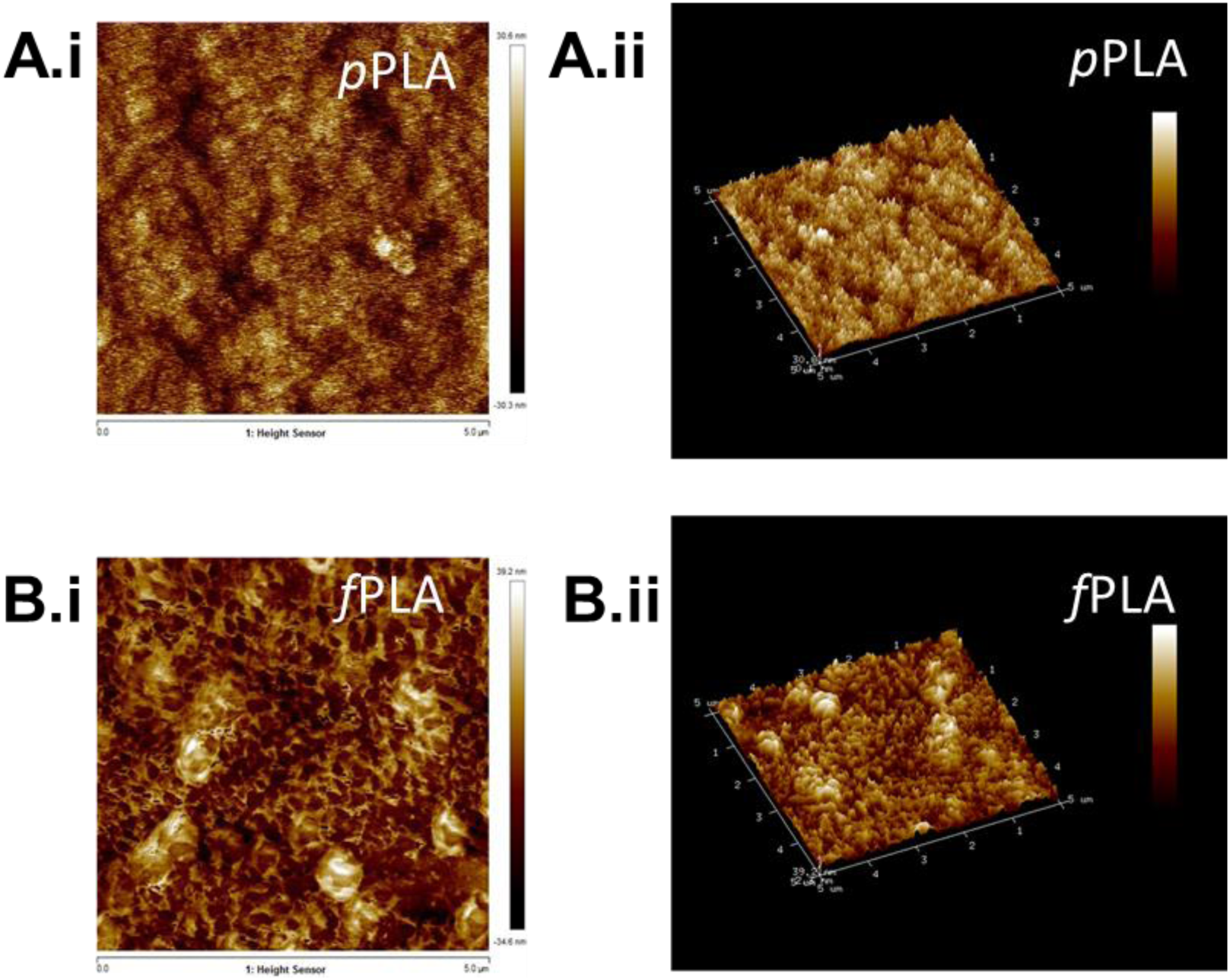
Atomic Force Microscope Imaging of Pristine and Functionalized PLA. Atomic force Microscope Imaging of Pristine and Functionalized PLA was performed with Dimension FastScan Atomic Force Microscope (ScanAsyst, Bruker), operating in tapping mode with a triangular Silicon Nitride cantilever (FASTSCAN-A probe, Bruker) at resonant frequency of ≈1500 kHz. The scan field was 5×5 µm^2^ and the amplitude of the height profile was recorded, with maximum peak values of 39.2 nm for the functionalized PLA (fPLA) corresponding to the rod formation already seen in the interferometry data, Figure 1.

**Figure SI2.**
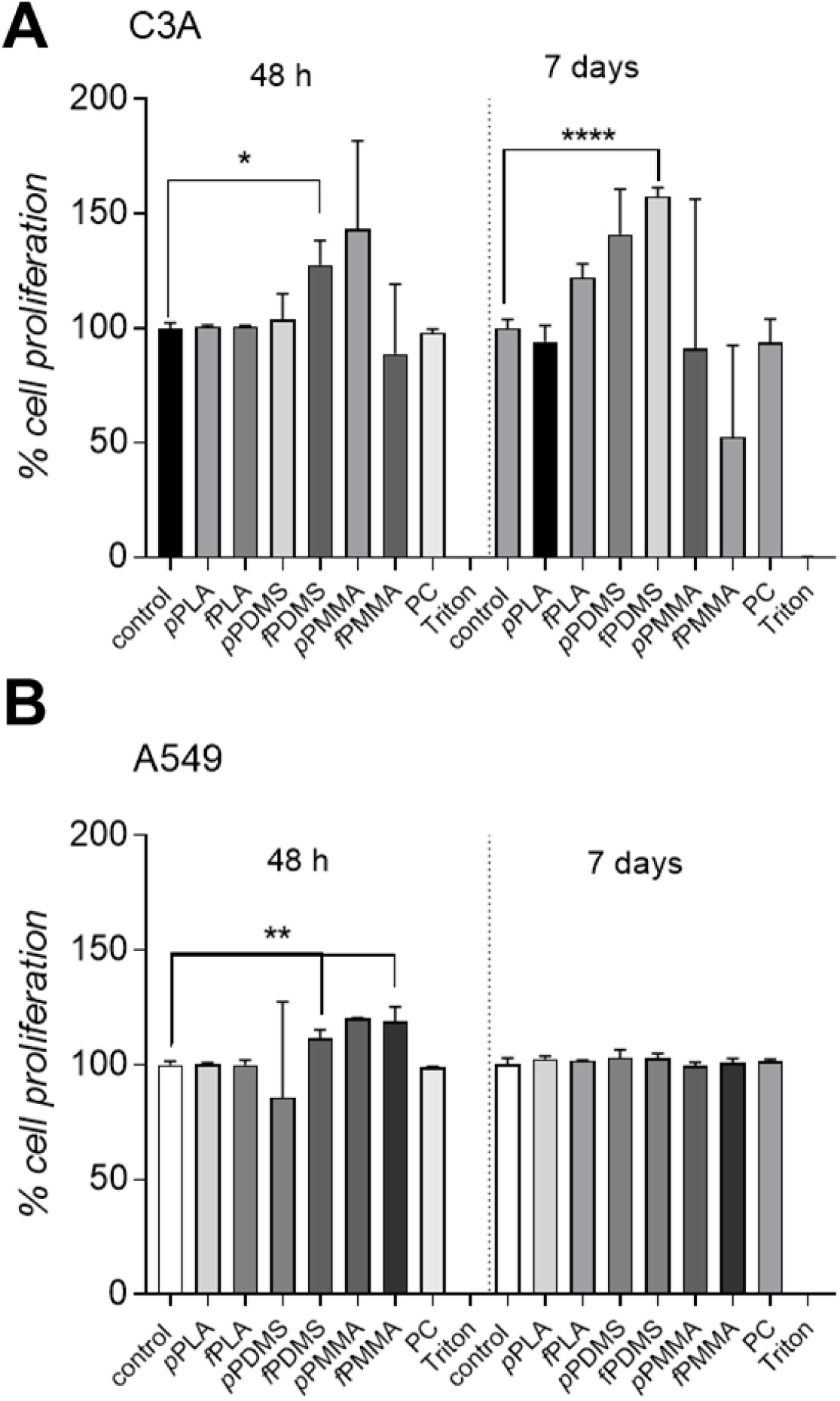
Cell Proliferation data for C3A (A) and A549 (B) on uncoated discs of fPLA,pPLA PDMS, fPDMS, fPMMA, pPMMA, PC. The positive control is an untreated PS well and the negative control is a untreated PS well with Triton.

**Figure SI3.**
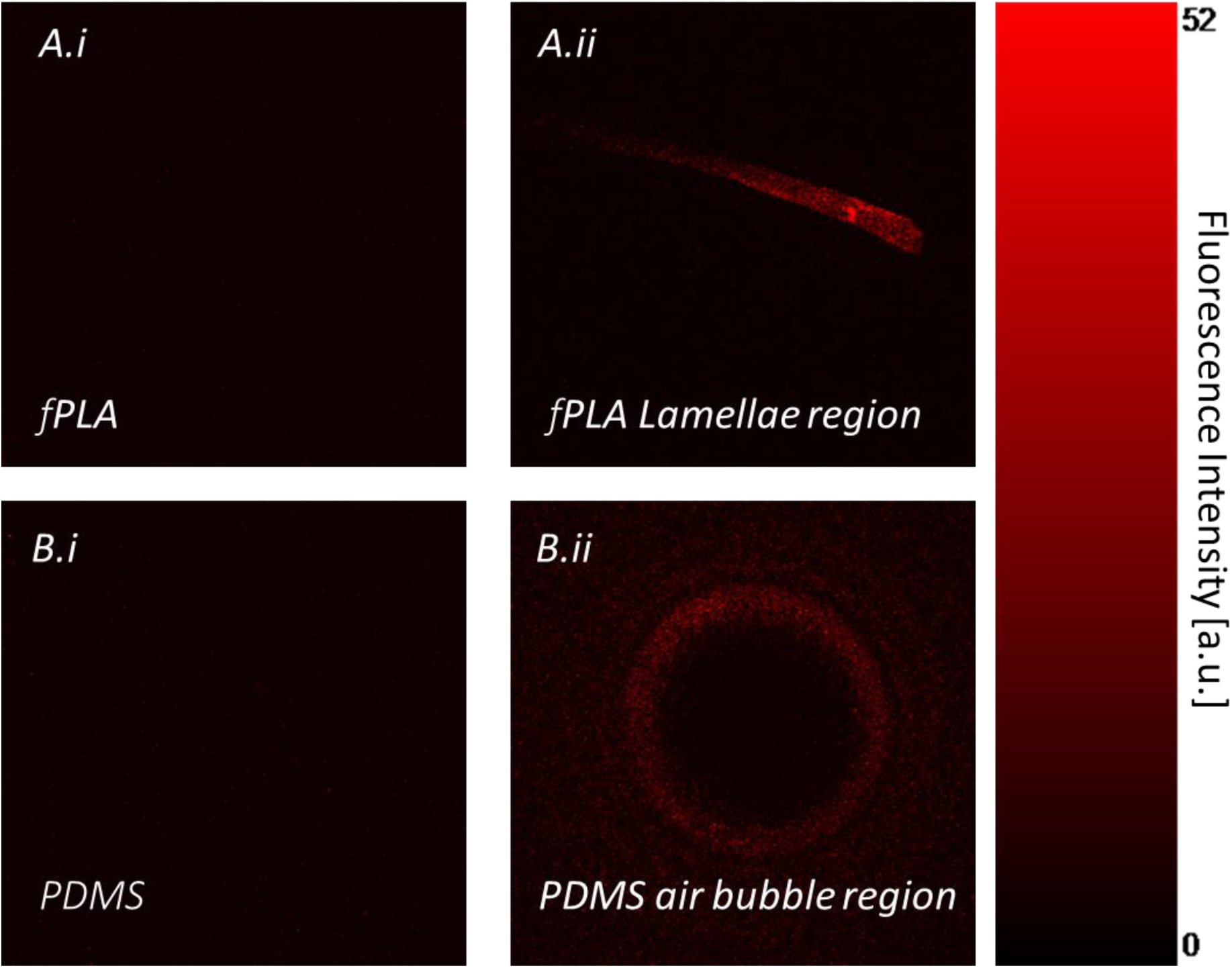
PLA Optical properties. Ai) Fluorescent background of well manufactured in 1 mm thick sample of fPLA. No background fluorescence is noticeable. A,ii) Fluorescent background of well manufactured in 1 mm thick sample of fPLA with a defect. An increased fluorescent background around a lamellae indicates a localized crystallisation. It is important that PLA pellets are not dried at temperature above 50°C (PLA glass transition temperature). Such defect indicates sub-optimal drying condition before sheet moulding B.ii) Fluorescent background of well manufactured in 1 mm thick sample of PDMS with some trapped bubble due to incorrect degassing process. An increased fluorescent background in visible around the bubble. The presence of an air bubble represents a discontinuity inside the bulk of the material which results in an increase auto-fluorescent background.

**Figure SI4.**
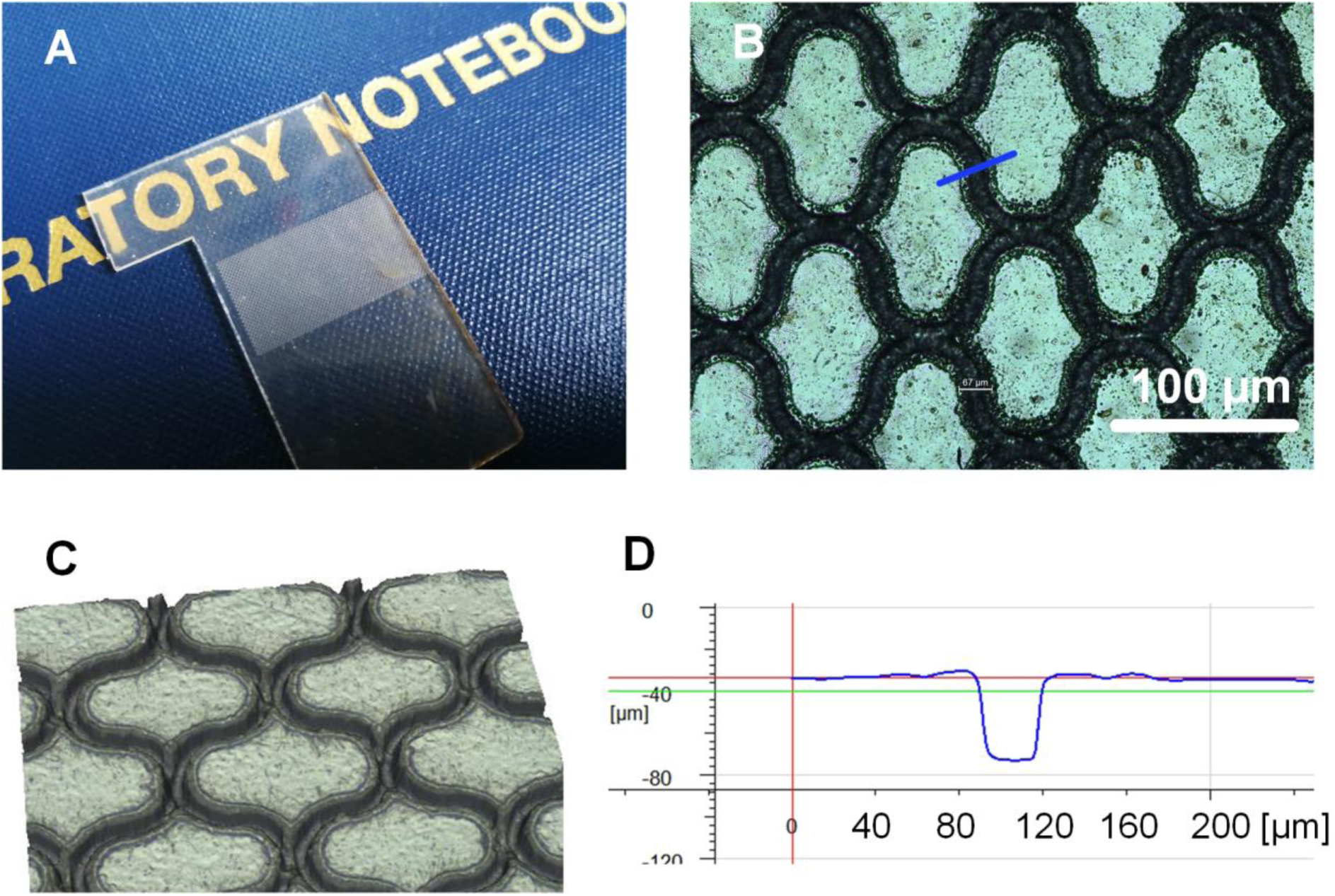
Production of microchannels with sub-100 µm width. A CO_2_ laser cutter enables limited resolution and minimum channel width. In our experience, regardless of the Epilog CO_2_ laser input parameters chosen (e.g. laser power, speed, frequency and focus), channel widths smaller than 100 µm are impossible to achieve with good reproducibility in PLA, or any other material. However, when channel widths with size between 20 – 100 µm are needed (e.g. for the production of perfusion barriers) picosecond lasers can be used. Here, as a proof of concept, a complex 2D structure has been generated on PLA using a picosecond laser micromachining system (Trumpf TruMicro 5×50) at Heriot-Watt University. The laser system enables the machining of various materials (including glass and transparent polymers) using a laser beam of the diameter smaller than 35μm (as measured at 1/e2 of its maximum intensity). A) photograph of the PLA sample with sub-100 µm honeycomb structures. B) The honeycomb structure was generated using a 24 μm diameter laser spot of the wavelength of 515 nm. The pulse energy was 46.9 μJ, pulse repetition frequency was 40 kHz, whereas the laser beam scan speed was 80 mm/s. The laser system allows the generation of microchannels on PLA with different depths and reproducible widths below 100 µm. C) Alicona surface profile showing the honeycomb channel features with depths of 40 µm D) Example surface profile obtained on the Alicona profilometer. The cross-section represented here is indicated in blue on B) and is measured to be 40 × 67 µm.

